# Base modification analysis in long-read sequencing data using minimod

**DOI:** 10.1101/2025.07.16.665072

**Authors:** Suneth Samarasinghe, Clare M. Robinson, Jonathan Göke, Ira W. Deveson, Hasindu Gamaarachchi

## Abstract

Recent advances in long read sequencing technologies have enabled the detection of various DNA and RNA base modifications in addition to standard nucleotide sequences. Both major vendors in this space, Oxford Nanopore Technologies (ONT) and Pacific Biosciences (PacBio), now include base modification information in their sequencing outputs using MM/ML tags embedded in unaligned BAM files. Each vendor also provides dedicated tools for extracting and analysing these tags, such as ONT’s modkit and PacBio’s pb-CpG-tools. This work presents minimod, a vendor-independent tool designed to extract and analyse base modifications from sequencing data generated by platforms that support MM/ML tags. Benchmarking revealed that for DNA data, minimod was ~1.3-1.6× faster on a server and ~4× on a laptop compared to modkit and pb-CpG-tools. For RNA data, minimod achieved even greater speedups compared to modkit, ~12× on the server and ~55× on the laptop. Minimod is a free, open-source application written in C and is available at https://github.com/warp9seq/minimod.

## 1. Introduction

Base modifications, which are chemical alterations of canonical nucleotide bases, are known to play a critical role in regulating gene expression across diverse cell types [1, 2]. They have been studied in multiple areas spanning from disease and ageing to forensic biology [3–5]. There are dozens of known DNA modification types [6] and well over a hundred types for RNA [7]. The most prevalent DNA base modification in mammals is 5-methylcytosine (5mC), which occurs predominantly at CpG dinucleotides and less frequently in non-CpG contexts [8, 9]. Methylation levels in the promoter region of a gene can regulate its expression by switching the gene on or off [8]. Such silencing of tumour suppressor genes is known to increase an individual’s risk of developing certain cancers [10].

Traditionally, bisulphite sequencing, typically performed using short-read sequencing platforms, has been used to identify DNA methylation [11]. Bisulphite conversion can cause complications due to extensive fragmentation of DNA during deamination, and reliance on short reads limits detection to only short-range methylation patterns [12]. More recently, long-read technologies such as Pacific Biosciences (PacBio) single-molecule real-time (SMRT) sequencing and Oxford Nanopore sequencing have become increasingly popular for detecting DNA methylation. Their substantially longer reads enable the analysis of long-range methylation patterns [12, 13]. In addition to 5mC, long-read sequencing technologies can detect various other DNA modifications, including 4mC, 5hmC, and 6mA [14–16]. ONT’s native direct RNA sequencing can detect RNA modifications such as m5C, m6A, inosine, pseU, and 2’-O-methyl [17–20].

Both major long-read vendors, ONT and PacBio, now produce unaligned SAM/BAM files that encode base modifications using auxiliary MM and ML tags, as defined by the official SAM tags specification. These tags can be retained during read alignment to a reference genome and subsequently used in downstream analyses, such as computing base modification frequencies. For such analyses, vendor-specific tools can be used, for example, PacBio’s pb-CpG-tools and ONT’s modkit. Although both tools are open-source, they are distributed under non-standard licenses: pb-CpG-tools under the Pacific Biosciences Software License Agreement, and modkit under the Oxford Nanopore Technologies PLC Public License 1.0. Such non-standard licenses often impose restrictions. For example, the Pacific Biosciences Software License Agreement limits the use of its software to data generated exclusively on PacBio instruments, while the Oxford Nanopore Technologies PLC Public License restricts commercial use.

Here, we introduce minimod, a fully open-source (MIT-licensed), vendor-independent software tool capable of processing base modifications from SAM/BAM files. Minimod supports platforms that encode modification information using MM/ML tags, including those from ONT and PacBio. Minimod supports both DNA and RNA modifications.

Our benchmarking revealed that minimod was ~1.3-1.6× faster on a server and ~4× faster on a laptop compared to both modkit and pb-CpG-tools when processing DNA data. For RNA data, minimod delivered a significant performance improvement, achieving ~12× faster execution on the server and ~55× on the laptop compared to modkit.

## 2. Methods and implementation

Minimod is implemented using the C programming language and provides two core subtools: *view* for extracting modifications along with modification probabilities; and, *freq* for computing modification frequencies. Both subtools take a BAM file containing modification tags and the reference genome as input.

### 2.1. Minimod software architecture

Minimod is designed and implemented for high performance and efficient resource utilisation. Minimod supports parallel processing through multithreading and pipeline stages implemented using POSIX threads. For a small memory footprint and overhead, hash tables are implemented using the lightweight *khash* from the klib library. htslib is used to load BAM files and to retrieve SAM auxiliary tags.

#### 2.1.1. Data structure initialisation

In the initialisation phase, command line arguments are parsed, validated and stored to be used in later steps. All possible data structures are allocated and initialised, including those used within threads. Such allocated data structures are reused later whenever possible to minimise the time overhead of dynamic memory allocations during runtime.

After memory allocation, the reference genome/transcriptome is loaded into memory. When loading the reference, we perform the Knuth-Morris-Pratt pattern searching algorithm on each reference sequence to identify the contexts requested by the user and mark them in a genome-wide array. This array is used later to identify if a modification lies within a requested context (section 2.2.4). The reference sequence is then unloaded to save memory.

#### 2.1.2. Parallel processing

After initialisation, the processing of BAM records is performed in batches (batch size can be user-specified, see section 2.2.3) using three interleaved pipeline stages. Both subtools have *load* and *process* as the first two pipeline stages. The third pipeline stage in the *view* subtool is *output*, while in the *freq* subtool it is *merge*.

##### *view* subtool

In the *load* pipeline stage, a batch of BAM records is loaded into memory from the disk. For decompressing BAM records, multiple threads are used. In the *process* pipeline stage, multiple threads are launched to compute base alignment and extract modifications from MM and ML tags (see section 2.1.3). These decoded modifications are stored in per-record data hash tables after checking if they lie within a requested context using the context array initialised in section 2.1.1. Finally, the *output* pipeline stage prints the extracted modifications to a given destination.

##### *freq* subtool

The *load* and *process* stages are identical to those in the *view* subtool above. When computing modification frequencies, we need to aggregate modifications based on contig name, position, modification type and strand (and haplotype if specified, see section 2.2.6), and compute a frequency value for each entry. This is achieved by merging modifications from per-record hash tables into a global hash table in the *merge* pipeline stage. This is performed by a single thread to avoid the need for locking. Before updating an entry in the hash table, a modification call is categorised based on the modification probability values as discussed in section 2.2.5. Fig. 1A shows how each batch progresses through the three pipeline stages of the *freq* subtool. The *load* stage of the second batch starts together with the start of the *process* stage of the first batch. Finally, when the last batch is merged, a single-threaded output stage writes the modifications along with frequency values.

**Fig 1:**
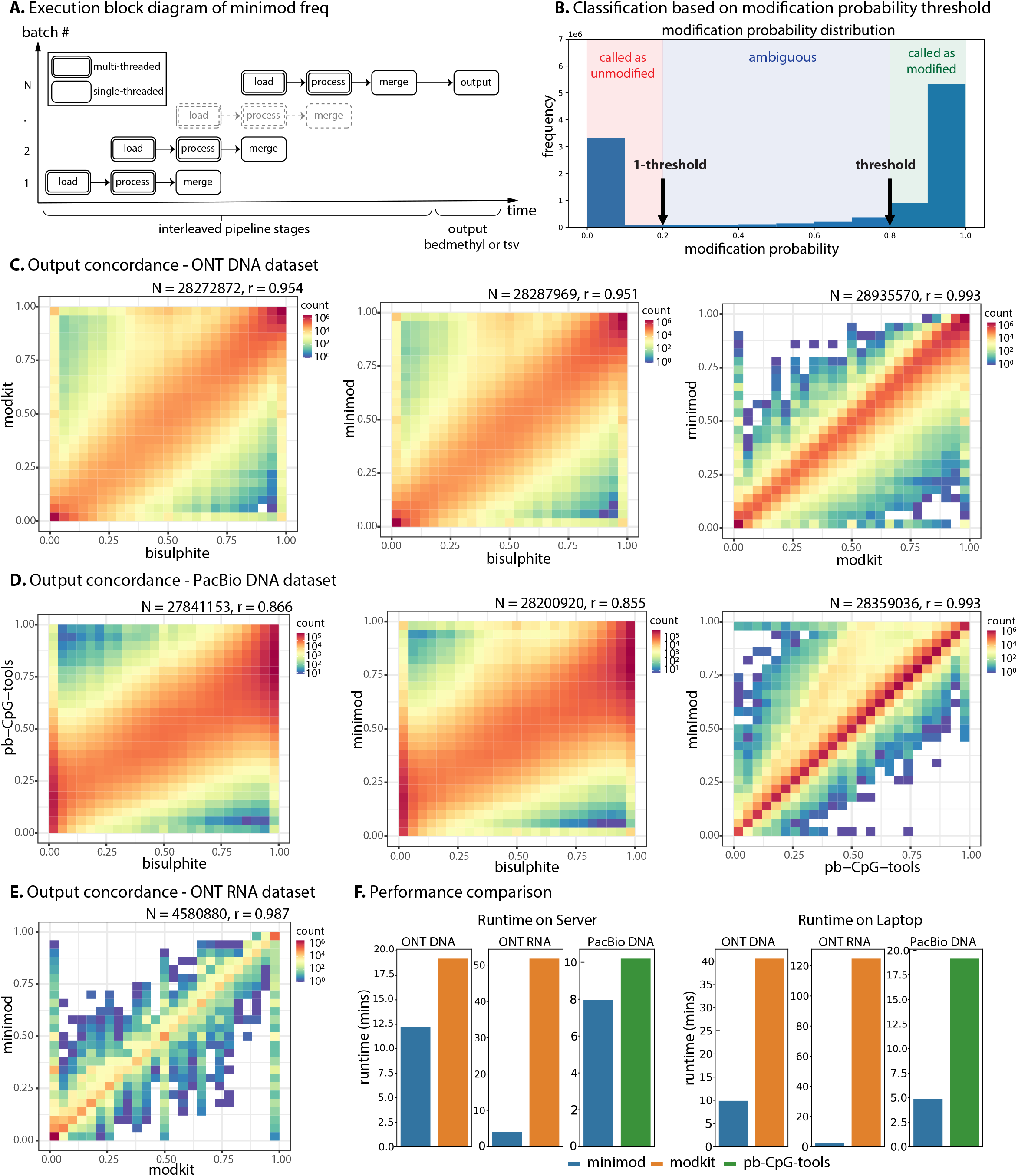
**A)** Execution block diagram of *minimod freq* with interleaved stages, load (load a batch of records from input BAM), process (extract modifications from the batch), merge (merge modifications into a genome-wide hash table), followed by the final non-interleaved output stage. **B)** Classification of the modifications based on the modification probability threshold. **C)** Correlation of methylation frequency results for the ONT DNA dataset. Panel 1 compares modkit with bisulphite, panel 2 compares minimod with bisulphite, and panel 3 compares minimod vs modkit. **D)** Correlation of methylation frequency results for the PacBio DNA dataset. Panel 1 compares pb-CpG-tools with bisulphite, panel 2 compares minimod with bisulphite, and panel 3 compares minimod vs pb-CpG-tools. **E)** Correlation of methylation frequency results for the ONT RNA dataset (aligned to transcriptome) with minimod vs modkit. **F)** Execution time of modification frequency computation using minimod, modkit and pb-CpG-tools on a server and laptop computer.

#### 2.1.3. BAM record processing

When computing base alignment, the aligned reference position is calculated by iterating through the CIGAR string of a BAM record. The read-to-reference alignment is stored in an array indexed by read position.

When extracting modifications from MM and ML tags, minimod iterates through and decodes the MM tag, which is a string that indicates the canonical base, strand, type of modification, and the location of modification as a list of delta-encoded positions. Similarly, minimod decodes the ML tag, which is a byte array containing modification likelihoods (probabilities) for each modification reported in the MM tag. The MM/ML tags are parsed using our efficient implementation described in realfreq [21].

### 2.2. Features and usage

The only external dependencies of minimod are *htslib* and *zlib*; thus, minimod can be compiled with minimal effort. Also, binaries for x86_64 are provided for further convenience of the user. Usage of currently available core subtools and their features are as follows.

#### 2.2.1. view subtool

The *view* subtool extracts modifications from auxiliary SAM tags and writes to the output in TSV format. A simple usage command is as follows, which writes all 5mC modifications found in the CpG sites (default modification code “*m*” and context “*CG*”, explained in section 2.2.4) to mods.tsv file.

~~~
minimod view ref. fa reads. bam > mods. tsv
~~~

#### 2.2.2. freq subtool

The *freq* subtool extracts modifications and aggregates them based on contig, position, and modification type to compute frequency values, followed by writing the output in TSV format (default) or BED format (if specified with -b flag). A simple example command is given below.

~~~
minimod freq ref. fa reads. bam > modfreqs. tsv
~~~

This writes the modification frequencies for 5mC found in CpG sites (default modification code “*m*” and context “*CG* “, explained in section 2.2.4), considering 0.8 as the default modification threshold (explained in section 2.2.5), to the modfreqs.tsv file.

#### 2.2.3. Batch size and threads

As discussed before, minimod processes batches of BAM records, with the *load* and *process* pipeline stages utilising multithreading. The batch size and the number of threads are user-configurable options, allowing adjustment to suit the available system memory and number of CPUs. The *-K* option sets the maximum number of BAM records per batch, *-B* option sets the maximum number of bases per batch and *-t* sets the number of threads.

#### 2.2.4. Filter by modification code and context

For both subtools *view* and *freq*, minimod can output a specific type of modification observed in a specific context in the reference using the *-c* option. It takes an argument in the format of *x* or *x* [*y*]. Here, *x* is a single character representing the modification type/code specified in the official SAM tags specification (e.g., *m, h*) or a ChEBI number for non-standard modifications. *y* is a multi-character context (e.g., *CG, A*, or ^*\**^ representing all contexts). If only *x* is specified, the default context is assumed for *y*. The following are a few example arguments for the *-c* option.

- *m*[*CG*] (or just *m*): 5mC modifications in CG context
- *h*[*CG*] (or just *h*): 5hmC modifications in CG context
- *a*[*A*] (or just *a*): m6A modifications of A bases
- *a*[^*\**^]: m6A modifications regardless of the base in the reference
- *17802* [*T*]: pseU (ChEBI: 17802) modifications in T context

Minimod by default uses the reference to precisely match the context when filtering modifications based on context (unless ^*\**^ is given as the context). This is because basecalling errors may be present in the reads, resulting in a read context that does not reflect the true sequence at that position.

#### 2.2.5. Filter by modification threshold

The modification threshold value is only applicable to the *freq* subtool and can be specified by the -m option. If not specified, the default threshold value of 0.8 is assumed. It is used to categorise modification calls as modified, unmodified, or ambiguous, based on the modification probability as follows. It is visually represented in Fig. 1B. Read calls that are categorised as ambiguous are excluded from the calculation of modification frequency (both within total_modified and total_called values).

~~~
if p(5 mC) >= 0.8 [threshold]:
   considered as called (5 mC) and modified (5 mC)
else if p(5 mC)<= 0.2 [1 - threshold]:
   considered as called (5 mC)
else :
   ignored as ambiguous
mod_freq (5 mC) = total_modified (5 mC)/ total_called (5 mC)
~~~

#### 2.2.6. Haplotypes and modifications in insertions

Minimod can output the haplotype information in a separate column for both *view* and *freq* subtools by specifying the *--haplotypes* flag. When enabled with *freq*, in addition to separate entries for each haplotype, an additional entry will be printed considering all haplotypes indicated by a ^*^ in the haplotype column. Furthermore, modifications within inserted regions can be included in the output of *view* subtool, by specifying the *--insertions* flag. This will add a new column, *ins_offset*, with the position of the modified base within the inserted region. However, the *--haplotypes* and *--insertions* flags are only applicable when the output is in TSV format.

## 3. Evaluations and results

### 3.1. Datasets and computer systems

To evaluate DNA modifications, we used two HG002 whole-genome datasets: a 151.8 Gbase dataset sequenced on an ONT PromethION (henceforth referred to as *ONT DNA*) and an 84.4 Gbase dataset sequenced on a PacBio Revio (*PacBio DNA*). To evaluate RNA modifications, we used a 17.8 Gbase dataset of a Universal Human Reference RNA sample sequenced using an ONT PromethION (*ONT RNA*). See Supplementary Table S1 for details of the datasets. HG002 Bisulphite data was downloaded from the publicly available ONT open-data AWS repository (s3://ont-opendata/gm24385_mod_2021.09/bisulphite/cpg).

We conducted benchmarks on a high-performance server and a laptop to evaluate minimod and compare its performance with ONT’s modkit and PacBio’s pb-CpG-tools. The server computer was equipped with two Intel Xeon Gold 6154 CPUs and 376 GB of RAM. The laptop computer used for the evaluation was equipped with an Intel i7-1370P CPU and 32 GB of RAM. Both systems were running Ubuntu 22.04 as the operating system. See Supplementary Table S2 for more details of the systems.

### 3.2. Evaluation

We evaluated minimod in terms of concordance with other tools and execution performance.

The ONT datasets were first converted from POD5 to BLOW5 using blue-crab [22, 23]. The ONT DNA dataset was basecalled using buttery-eel [24] with *5hmc_5mc* modification calling in *CG* context. The ONT RNA dataset was basecalled using slow5-dorado [25] with *m6A_DRACH* modification calling. This produced unaligned SAM/BAM output with MM/ML tags. The PacBio DNA dataset was already in the unaligned SAM/BAM format with MM/ML tags. The unaligned SAM/BAM files were aligned using minimap2 [26] (with *-Y -y* options to retain MM/ML tags) and were sorted using samtools [27] to produce BAM files with MM/ML tags. As the reference, the hg38 genome and the Gencode v40 transcriptome were used.

To measure the concordance, modification frequency was obtained by running minimod *freq* on all three datasets (sorted BAM with MM/ML tags), modkit *pileup* on two ONT datasets, and pb-CpG-tools on the PacBio dataset. Methylation (5mC) modification was considered for the two DNA datasets and N6-methyladenosine (m6A) for the

RNA dataset. Comparison of methylation frequency results and plotting the correlation were performed by adapting the scripts *compare_methylation*.*py* and *plot_methylation*.*R* from the f5c/nanopolish repository [28, 29] to support the minimod TSV format.

To measure execution performance, each experiment was performed five times on both laptop and server computers. We used */usr/bin/time* with *-v* flag when running each tool and took elapsed (wall clock) time as the runtime metric for performance evaluation and maximum resident set size as the peak RAM consumption. All software was executed with 32 threads on the server and with 8 threads on the laptop.

Unless otherwise stated, minimod v0.4.0, modkit v0.5.0 and pb-CpG-tools v2.3.2 were used (latest available at the time of benchmarking). All software versions and full commands that we executed are in Supplementary Note 1.

### 3.3. Output concordance

For the ONT DNA dataset, both modkit and minimod showed similar Pearson correlation coefficients of ~0.95 when compared against the bisulphite truthset (Fig. 1C, first and second panels). However, minimod had ~15,000 more genomic positions shared with the truthset than modkit. When compared with each other, minimod vs modkit had a correlation of ~0.99 (Fig. 1C, third panel).

For the PacBio DNA dataset, methylation results from pb-CpG-tools showed a slightly higher correlation (~0.87) than minimod (~0.86), as shown in Fig. 1D (first and second panels). However, minimod had ~360,000 more data points in common with the truthset than pb-CpG-tools. When compared with each other, minimod vs pb-CpG-tools showed a correlation of ~0.99 (Fig. 1D, third panel).

For the ONT RNA dataset, m6A modification results between modkit vs minimod showed a correlation of ~0.99 when aligned to the transcriptome (Fig. 1E). We added support for RNA alignments to the genome in minimod v0.5.0, which also showed a ~0.99 correlation with modkit (Supplementary Fig. S1).

The differences in the number of data points reported by each tool, as well as the small differences in correlation, arise from differences in the default filtering methods and threshold values used by modkit and pb-CpG-tools relative to minimod (section 2.2.5 and Supplementary Note 2). When configured with matched settings and thresholds, minimod and modkit produce identical results for two-class modification models, as demonstrated in Supplementary Note 2. For models with three or more classes, negligible discrepancies may remain between the two tools even with matched thresholds (correlation 0.99959, Supplementary Note 3). This is because minimod classifies each modification code independently against its own probability, whereas modkit discards modifications whose probability falls below its threshold (including canonical class) and assigns a most likely call (Supplementary Note 3). Note that output from minimod *view* and modkit *extract* remain identical, as thresholding is not applied in these modes, unlike during frequency computation.

### 3.4. Performance comparison

In our benchmarks, minimod was faster than the corresponding vendor tool for all three datasets on both the server (Fig. 1F left panel) and the laptop (Fig. 1F right panel) when processing modification frequencies. On the server, minimod was ~1.6× faster than modkit when processing the ONT DNA dataset, ~12.6× faster than modkit when processing the ONT RNA dataset (transcriptome alignment), and ~1.3× faster than pb-CpG-tools when processing the PacBio DNA dataset. On the laptop, minimod was ~4.1× faster than modkit when processing the ONT DNA dataset, ~54.8× faster than modkit when processing the ONT RNA dataset (transcriptome alignment), and ~4.0× faster than pb-CpG-tools when processing the PacBio DNA dataset.

The substantial performance advantage of minimod over modkit for RNA was observed regardless of whether reads are aligned to the transcriptome or the genome. To confirm this, we repeated the RNA experiment using reads aligned to the hg38 genome on the server computer, using minimod v0.5.0, which introduced support for RNA-to-genome alignments. Minimod took only 5.3 minutes compared to 1.1 hours for modkit, representing an ~12.5× performance speedup for minimod, which is consistent with the performance improvement observed on the transcriptome.

Minimod maintained a memory footprint of less than 16 GB across all datasets and computing platforms. On the server computer, minimod used 10.8 GB of memory when processing the ONT DNA dataset, comparable to modkit which used 11.1 GB (Supplementary Fig. S2). For the ONT RNA dataset, minimod used a higher amount of memory (8.5 GB for transcriptome, 15.7 GB for genome) compared to modkit (6.6 GB for transcriptome and 3.0 GB for genome). On the PacBio DNA data, minimod used 13.8 GB compared to 4.8 GB by pb-CpG-tools. On the laptop, minimod used 10.5 GB of memory for ONT DNA, 7.2 GB and 14.4 GB for ONT RNA (transcriptome and genome, respectively), and 10.8 GB for PacBio DNA. While minimod uses more RAM than modkit and pb-CpG-tools (Supplementary Fig. S2), its memory footprint can be reduced to ~9 GB for the hg38 genome and ~2 GB for the Gencode v40 transcriptome by decreasing the batch size using the -K and -B options.

## 5. Authors’ contributions

H.G. and I.W.D. conceived the project. H.G. and S.S. designed the software with contributions from all authors. S.S. implemented the software. H.G., J.G. and I.W.D. designed the experiments. S.S. and C.M.R. performed benchmarking experiments, data analysis and validation. S.S. and H.G. prepared the manuscript with input from all authors. C.M.R. and J.G. provided expertise on modification analysis and critically revised the manuscript.

## 6. Acknowledgements

We thank our colleagues James Ferguson, Leah Kemp, Marjan Naeini, Jillian Hammond, Igor Stevanovski, Tonia Russell and Andre Martins Reis at the Garvan Institute for their assistance at different stages of this work.

## 8. Data and code availability

The source code for minimod is open-source at https://github.com/warp9seq/minimod under the MIT license. Minimod v0.4.0 is also available at doi:10.5281/zenodo.15876999. Unaligned and aligned BAM files with modification calls are available at doi:10.5061/dryad.c59zw3rm7. Raw nanopore signal data is available at ENA (HG002 sample: ERR12997168 and UHRR sample: ERR12997170).

## Supplementary Material

**Supplementary Table S1:** Datasets used for testing.

| Dataset name | Sample | Instrument | Sequencing details | Reads (millions) | Total seq. (Gbases) |
| --- | --- | --- | --- | --- | --- |
| ONT DNA | HG002 | PromethION | LSK114 kit, R10.4.1 pore, 5kHz data rate | 21.8 | 151.8 |
| ONT RNA | UHRR | PromethION | RNA004 kit, RP4 pore, 4kHz data rate | 16.5 | 17.8 |
| PacBio DNA | HG002 | PacBio Revio | 102-739-100 bindingkit, 102-118-800 sequencingkit, 102-202-200 smrtcellkit | 6.3 | 84.4 |

**Supplementary Table S2:** Computer systems used for testing.

| System | Description | n x CPU | total cores, CPU threads | RAM | Disk | O/S |
| --- | --- | --- | --- | --- | --- | --- |
| server | PowerEdge R740xd | 2 × Intel Xeon Gold 6154 | 36,72 | 376 GB | 12×10TB HDD drives(RAID6, ext4) | Ubuntu 22.04.5 LTS |
| laptop | Dell Precision 3480 | 13th Gen Intel Core i7-1370P | 14,20 | 32 GB | 1 TB NVMe SSD drive (NTFS) | Ubuntu 22.04.4 LTS |

**Supplementary Fig. S1.**
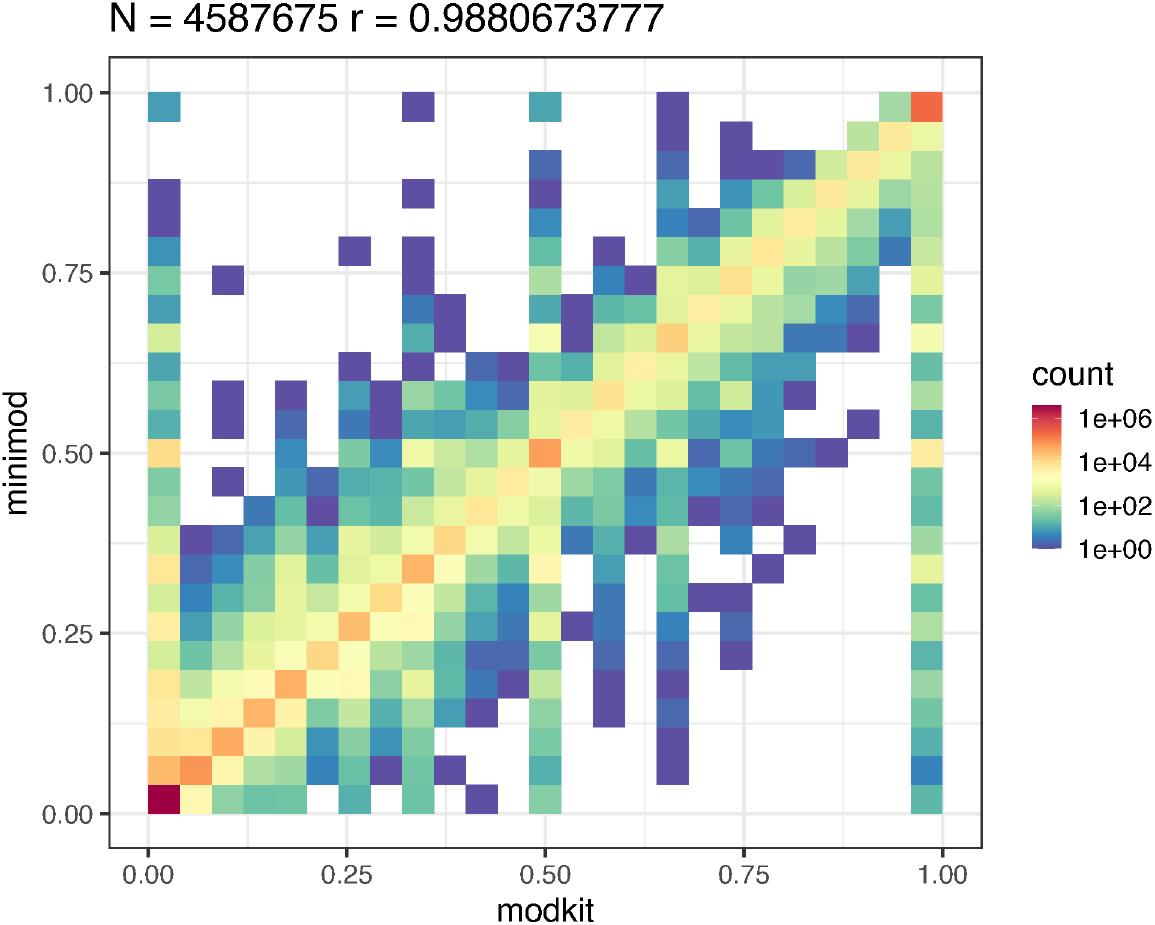
Correlation of methylation frequency results for the ONT RNA dataset (aligned to genome) with minimod vs modkit.

**Supplementary Fig. S2.**
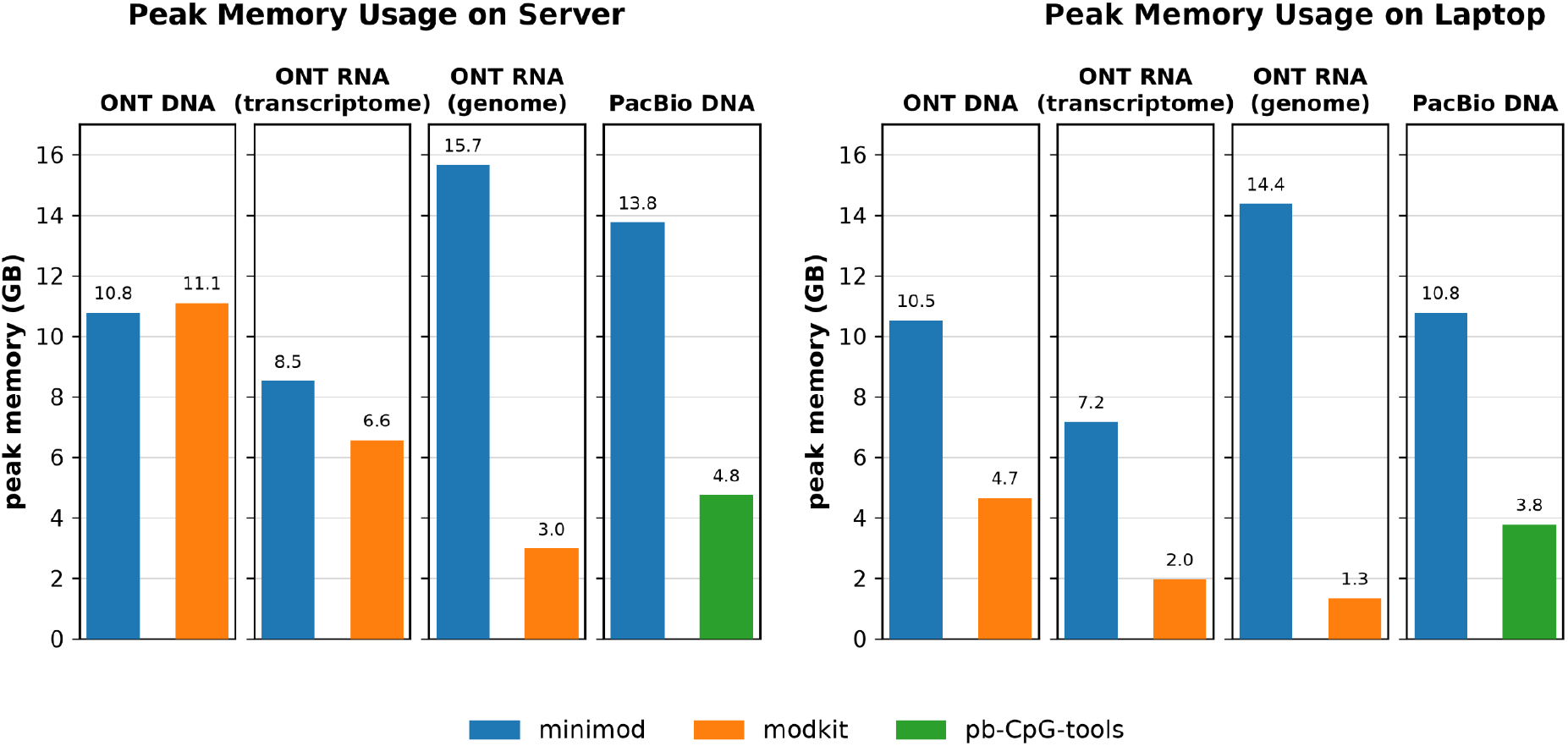
Peak RAM usage for modification frequency computation using minimod, modkit and pb-CpG-tools on server and laptop.

### Supplementary Note 1: Commands and versions of software used

#### Methylation analysis for ONT DNA

~~~
blue-crab p2s reads.pod5 -o reads.blow5
buttery-eel -g /dorado/bin/ --port 5000 --use_tcp --device cuda:all --call_mods –config
dna_r10.4.1_e8.2_400bps_5khz_modbases_5hmc_5mc_cg_hac.cfg -i reads.blow5 -o unaligned.sam
samtools fastq -TMM,ML unaligned.sam | minimap2 -ax map-ont -Y -y --secondary=no hg38noAlt.fa - |
samtools sort - -o sorted.bam && samtools index sorted.bam
# on server: THREADS=32, BATCH_K=3072, BATCH_B=40M
# on laptop: THREADS=8, BATCH_K=2560, BATCH_B=35M
/usr/bin/time -v modkit pileup --cpg --ref hg38noAlt.fa --ignore h -t ${THREADS} sorted.bam
modkit.bedmethyl
/usr/bin/time -v minimod freq -t ${THREADS} -K ${BATCH_K} -B ${BATCH_B} hg38noAlt.fa sorted.bam >
minimod.tsv
~~~

##### Versions

- buttery-eel: 0.6.0; ont-dorado-server: 7.4.12
- samtools: 1.21; minimap2: 2.28
- modkit: 0.5.0
- minimod: 0.4.0

#### Methylation analysis for PacBio DNA

~~~
samtools fastq -TMM,ML reads.bam | minimap2 -ax map-hifi -y -Y hg38noAlt.fa --secondary=no - |
samtools sort - -o sorted.bam && samtools index sorted.bam
# on server: THREADS=32, BATCH_K=5120, BATCH_B=105M
# on laptop: THREADS=8, BATCH_K=2048, BATCH_B=40M
/usr/bin/time -v pb-CpG-tools/bin/aligned_bam_to_cpg_scores --ref hg38noAlt.fa –bam sorted.bam
--output-prefix out/ --pileup-mode count --threads ${THREADS}
/usr/bin/time -v minimod freq -m 0.5 -t ${THREADS} -K ${BATCH_K} -B ${BATCH_B} hg38noAlt.fa sorted.bam
-o minimod.tsv
~~~

##### Versions

- samtools: 1.21; minimap2: 2.28
- pb-CpG-tools: 2.3.2
- minimod: 0.4.0

#### ONT RNA modification calling

~~~
blue-crab p2s reads.pod5 -o reads.blow5
slow5-dorado basecaller rna004_130bps_sup@v5.0.0 -x cuda:all reads.blow5 --modified-bases m6A_DRACH >
reads.unaln.bam
~~~

##### Versions

- slow5-dorado: 0.8.3

#### m6A analysis for ONT RNA (alignment to transcriptome)

~~~
samtools fastq -TMM,ML reads.unaln.bam | minimap2 -ax map-ont -Y -y --secondary=no -uf
gencode.v40.transcripts.fa - | samtools sort -o sorted.bam && samtools index sorted.bam
# on server: THREADS=32, BATCH_K=65536, BATCH_B=150M
# on laptop: THREADS=8, BATCH_K=57344, BATCH_B=120M
/usr/bin/time -v modkit pileup --ref gencode.v40.transcripts.fa -t ${THREADS} sorted.bam
modkit.bedmethyl
/usr/bin/time -v minimod freq -t ${THREADS} -K ${BATCH_K} -B ${BATCH_B} -c a
gencode.v40.transcripts.fa sorted.bam
-o minimod.mm.tsv
~~~

##### Versions

- samtools: 1.21; minimap2: 2.28
- modkit: 0.5.0
- minimod: 0.4.0

#### m6A analysis for ONT RNA (alignment to genome)

~~~
samtools fastq -TMM,ML reads.unaln.bam | minimap2 -ax splice -Y -y --secondary=no -uf hg38noAlt.fa - |
samtools sort -o sorted.bam && samtools index sorted.bam
# on server: THREADS=32, BATCH_K=65536, BATCH_B=150M
/usr/bin/time -v modkit pileup --ref hg38noAlt.fa -t ${THREADS} sorted.bam modkit.bedmethyl
/usr/bin/time -v minimod freq -t ${THREADS} -K ${BATCH_K} -B ${BATCH_B} -c a hg38noAlt.fa sorted.bam
-o minimod.mm.tsv
~~~

##### Versions

- samtools: 1.21; minimap2: 2.28
- modkit: 0.5.0
- minimod: 0.5.0

### Supplementary Note 2: Minimod output concordance with modkit in two-class classification

#### Differences in default parameters and settings

Due to the differences described below, minimod and modkit do not produce identical outputs with their default options. However, matching their parameters enables both tools to produce identical outputs for two-class modification models as demonstrated in the example walkthrough in the next section.

##### Relevant to minimod view and freq

- By default, minimod outputs *m* modified bases in the *CG* context that are mapped to and match the corresponding reference bases, unless a different modified base and/or context is specified with -c. In contrast, modkit outputs all modifications by default without requiring the modified base to match the corresponding reference base, unless a context is specified using *--cpg* or *--motif*.
- By default, minimod ignores secondary alignments and processes primary and supplementary alignments. Secondary alignments can be included using *--allow-secondary*. Minimod does not require an MN tag to process non-primary alignments. However, it terminates with an error when it detects hard clipping. By default, modkit ignores all non-primary alignments. The --allow-non-primary option can enable processing of secondary and supplementary alignments when a valid MN tag is present, as described [here](https://github.com/nanoporetech/modkit/blob/481e3c9e7930f3f499eadf1ef441606f33e6881c/book/src/intro_extract.md#note-on-non-primary-alignments).

##### Relevant to minimod freq only

- By default, minimod freq uses a modification threshold of 0.8 to determine whether a base is modified. In contrast, modkit pileup estimates the threshold programmatically from the data unless the user specifies one.
- Minimod freq applies the modification-filtering method described [here](https://github.com/warp9seq/minimod/blob/main/README.md#modification-threshold). In contrast, modkit pileup uses two-class and three-class modified-base calls, as described [here](https://github.com/nanoporetech/modkit/blob/v0.5.0/book/src/filtering_details.md).

#### Parameters and settings to get identical outputs from Minimod and Modkit

Now we demonstrate how one can obtain the identical output using minimod and modkit. We will use minimod v0.5.0 and modkit 0.5.0. The example below uses an RNA dataset mod+basecalled using slow5-dorado 1.1.1 (rna004_130bps_sup@v5.2.0, m6A_DRACH model) and aligned to the reference genome.

##### Dataset preparation

~~~
wget -O PNXRXX240011_reads_20k.blow5 https://slow5.bioinf.science/uhr_prom_subsubsample
slow5-dorado-1.1.1 basecaller rna004_130bps_sup@v5.2.0 PNXRXX240011_reads_20k.blow5
--modified-bases m6A_DRACH > rna_unaligned.bam
samtools fastq -T “MM,ML” rna_unaligned.bam | minimap2 -ax splice -uf hg38noAlt.fa
--secondary=no -t16 -y -Y - | samtools sort - -o rna.bam
samtools index rna.bam
~~~

##### Minimod freq vs Modkit pileup

**Frequencies of type “a” modifications in “A” context in RNA dataset:**

~~~
minimod freq -b -c a --skip-supplementary hg38noAlt.fa rna.bam > minimod.bedmethyl
modkit pileup --motif A 0 --filter-threshold A:0.8 --reference hg38noAlt.fa rna.bam
modkit.bedmethyl
test/compare.py modkit.bedmethyl minimod.bedmethyl
# Pearson correlation coefficient: 1.0
~~~

test/compare.py: Python script computes the Pearson correlation coefficient between two BED files. *compare*.*py file1*.*bedmethyl file2*.*bedmethyl* outputs the Pearson correlation coefficient of modification frequencies in two bedmethyl files.

**Supplementary Fig. S3.**
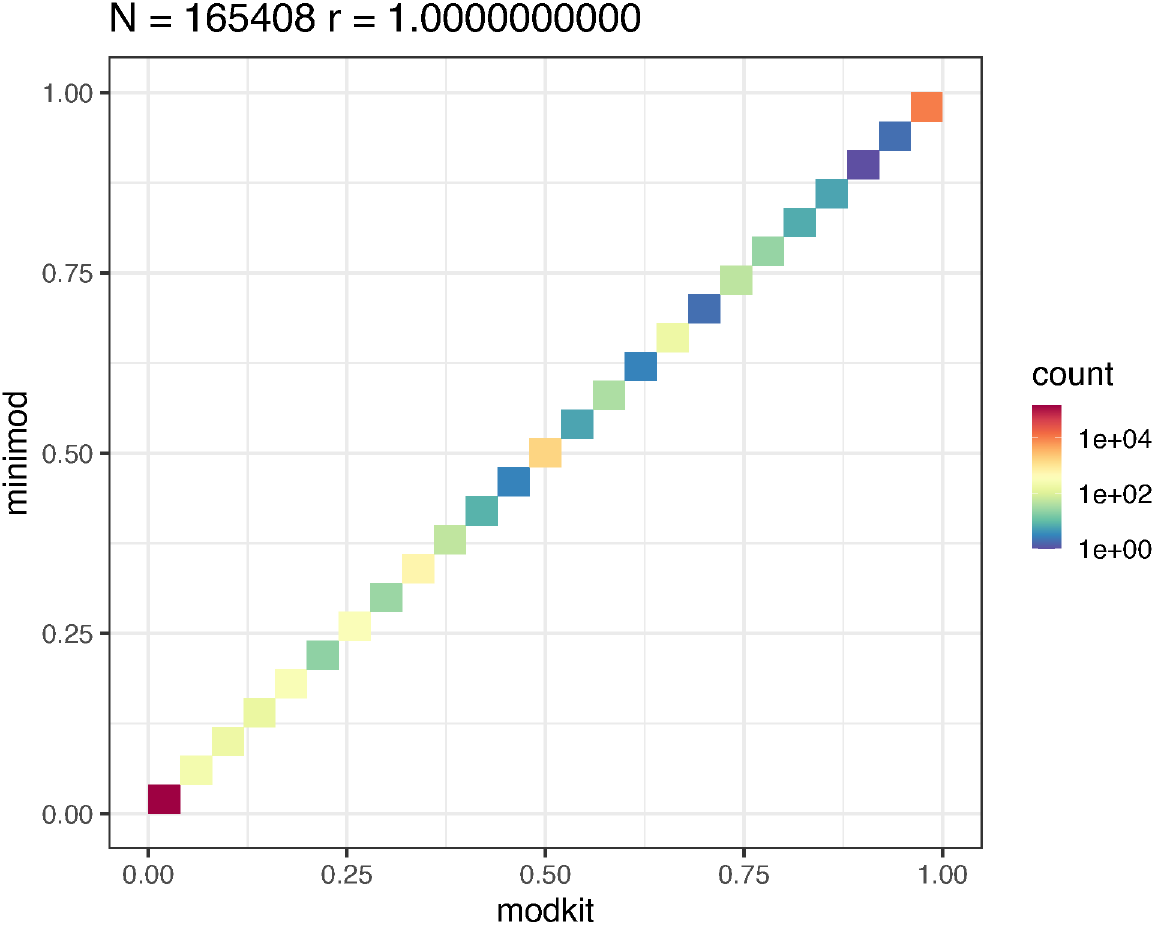
Correlation of methylation frequency results for the ONT RNA 20k dataset with minimod vs modkit.

###### Summary

When mod+basecalled using a 2-class classification model such as m6A_DRACH (classifies into modified as m6A or canonical A), minimod’s and modkit’s filtering methods result in the same outputs if the modification threshold is matched using -m in minimod or --filter-threshold in modkit:

~~~
minimod freq --skip-supplementary -m 0.9 -b -c “a[A]” hg38noAlt.fa rna.bam >
minimod.bedmethyl
modkit pileup --filter-threshold A:0.9 --motif A 0 --reference hg38noAlt.fa rna.bam
modkit.bedmethyl
test/compare_freq_bed_bed.sh minimod.bedmethyl modkit.bedmethyl compare_out
~~~

###### Results

~~~
compare_out/
in_both.tsv          # 162293 entries
large_freq_diff.tsv  # 0 entries
missing_in_file1.tsv # 0 entries
missing_in_file2.tsv # 0 entries
~~~

##### Minimod view vs Modkit extract full

###### View type “a” modifications in “A” context

~~~
minimod view -c ‘a[A]’ --skip-supplementary hg38noAlt.fa rna.bam > minimod.tsv
modkit extract full --motif A 0 --mapped-only --reference hg38noAlt.fa rna.bam modkit.tsv
test/compare_view_mktsv_mmtsv.sh modkit.tsv minimod.tsv compare_out
~~~

Alternative way to filter context from modkit output using awk instead of using --motif A 0, if modkit crashes due to out-of-memory:

~~~
modkit extract full --mapped-only --kmer-size 1 --reference hg38noAlt.fa rna.bam modkit.tsv
awk -F ‘\t’ ‘function c(b){b=toupper(b);return
b==“A”?”T”:b==“T”?”A”:b==“C”?”G”:b==“G”?”C”:b} NR==1 || ( $21==16 ? c($16)==“A” &&
toupper($16)==c($17) : toupper($16)==“A” && toupper($16)==toupper($17) )’ modkit.tsv >
modkit_A.tsv
test/compare_view_mktsv_mmtsv.sh modkit_A.tsv minimod.tsv compare_out
~~~

test/compare_view_mktsv_mmtsv.sh takes modkit’s extract tsv and minimod’s view tsv and creates four files in the out_dir:

- in_both.tsv : found in both files and probability difference is <= 0.002
- large_prob_diff.tsv : found in both files and probability difference is > 0.002
- missing_in_file1.tsv : missing in file1 which is modkit_extract.tsv
- missing_in_file2.tsv : missing in file2 which is minimod_view.tsv

There are similar scripts we provide for view comparison between different file formats:

- test/compare_view_mktsv_mmtsv.sh
- test/compare_view_mktsv_mktsv.sh
- test/compare_view_mmtsv_mmtsv.sh

###### Results

~~~
compare_out/
in_both.tsv          # 394652 entries
large_prob_diff.tsv  # 0 entries
missing_in_file1.tsv # 0 entries
missing_in_file2.tsv # 0 entries
~~~

### Supplementary Note 3: Minimod output concordance with modkit for multi-class classification

For models with three or more classes, the outputs from minimod and modkit converge closely but are not identical, for the reasons given below:

- minimod evaluates each modification code requested with -c independently. For a code with a modification probability of *p* and a threshold of *t*, a call is counted as modified if *p >= t*, as unmodified if *p <= 1* − *t*, and is discarded as ambiguous otherwise. Note that “unmodified” means not modified with respect to the modification type being analysed, and does not imply a canonical base. The canonical probability is never computed, and each modification code accumulates its own entry in the frequency output.
- modkit makes a single call per base. It first discards every class whose probability falls below its threshold, including the canonical class, whose probability is taken as *1-sum(p(mod))* and then filters each class (mods or canonical) as described in their documentation [here](https://github.com/nanoporetech/modkit/blob/v0.5.0/book/src/filtering_details.md).

For a two-class model, the above two methods are equivalent because, with a single modification code, p(canonical) = 1 − p(mod), so minimod’s test p <= 1 − t is the same test as modkit’s p(canonical) >= t, and the two outputs match once thresholds are matched. For three-class or higher models, the two methods are very close, but not equivalent:

- A base that modkit discards may be counted by minimod. Example: With threshold 0.8 and p(m6A) = 0.15, p(m1A) = 0.75, p(A) = 0.10, no class reaches 0.8, so modkit reports nothing. minimod counts the base as an unmodified call in the m6A entry (0.15 ≤ 0.2) and discards it as ambiguous in the m1A entry (0.2 < 0.75 < 0.8).
- One base can be counted as modified by more than one code. Example: If the threshold is set to a very low value like 0.4 and p(m6A) = 0.45, p(m1A) = 0.45, p(A) = 0.10, both codes satisfy p >= 0.4, so minimod counts the base as modified in the m6A entry and in the m1A entry. modkit may only report the most likely modification as per the documentation [here](https://github.com/nanoporetech/modkit/blob/v0.5.0/book/src/filtering_details.md). Note that a threshold of 0.4 reflects relatively low confidence for modified-site predictions, making it unlikely that users would apply such a permissive threshold in practice.

Having said that, if a sensible threshold like 0.8 is selected, the results will be very similar. Now we demonstrate this using an example. We will use minimod v0.5.0 and modkit 0.5.0. The example below uses a DNA dataset mod+basecalled using slow5-dorado 1.1.1 (dna_r10.4.1_e8.2_400bps_sup@v5.2.0, 5mCG_5hmCG model) and aligned to the reference genome.

#### Dataset preparation

~~~
wget -O PGXXXX230339_reads_20k.blow5
https://slow5.bioinf.science/hg2_prom_5khz_subsubsample
slow5-dorado-1.1.1 basecaller dna_r10.4.1_e8.2_400bps_sup@v5.2.0
PGXXXX230339_reads_20k.blow5 --modified-bases 5mCG_5hmCG > dna_unaligned.bam
samtools fastq -T “MM,ML” dna_unaligned.bam | minimap2 -ax map-ont hg38noAlt.fa
--secondary=no -t16 -y -Y - | samtools sort - -o dna.bam
samtools index dna.bam
~~~

**Supplementary Fig. S4.**
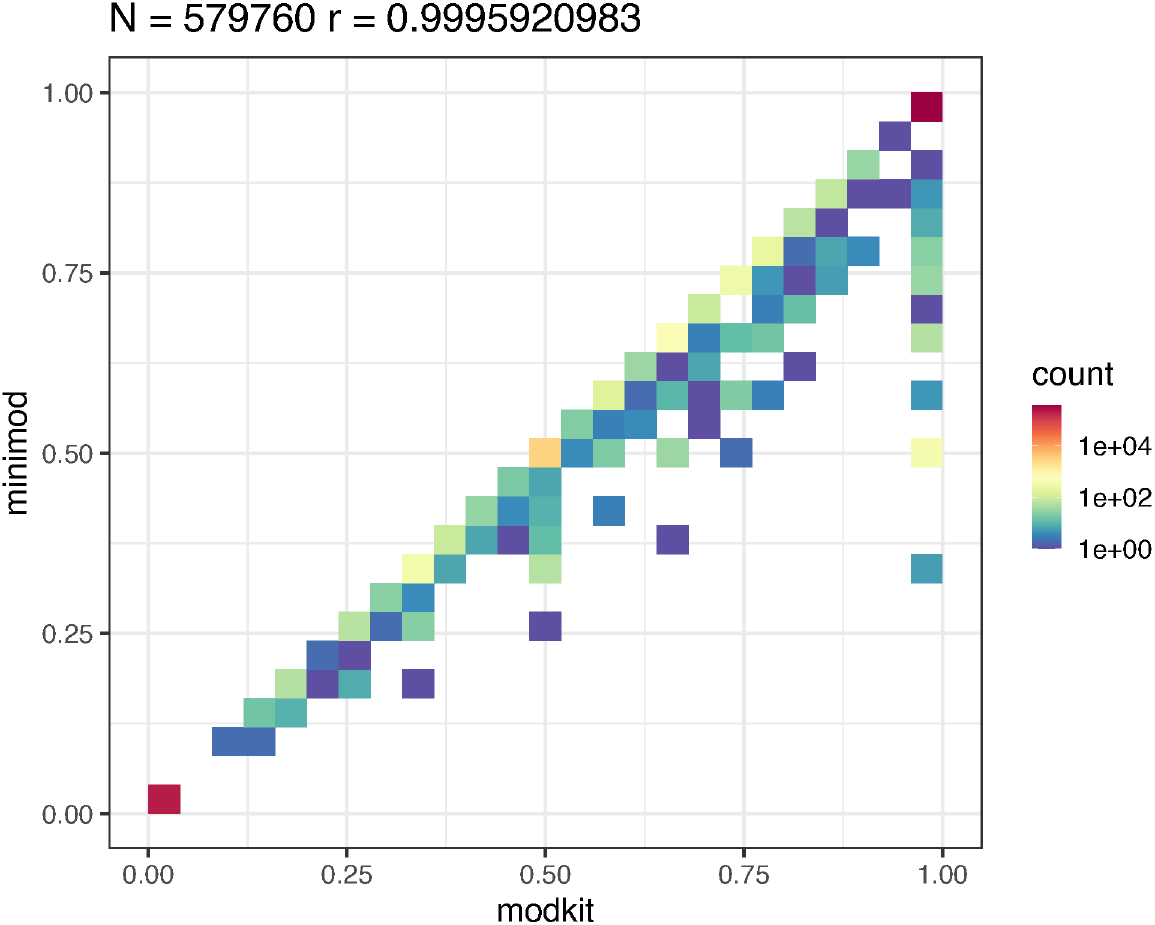
Correlation of methylation frequency results for the DNA 20k dataset with minimod vs modkit.

#### Minimod freq vs Modkit pileup

The following pair of commands using minimod v0.5.0 and modkit 0.5.0 should give similar outputs.

**Frequencies of type “m” modifications in “CG” context in DNA dataset:**

~~~
minimod freq -b --skip-supplementary hg38noAlt.fa dna.bam > minimod.bedmethyl
modkit pileup --cpg --filter-threshold C:0.8 --reference hg38noAlt.fa dna.bam modkit.bedmethyl
awk ‘$4==“m”’ modkit.bedmethyl > modkit_m.bedmethyl
test/compare.py modkit_m.bedmethyl minimod.bedmethyl
# Pearson correlation coefficient: 0.9995920972597717
~~~

#### Minimod view vs Modkit extract full

The following pair of commands using minimod v0.5.0 and modkit 0.5.0 should give identical outputs.

**View type “m” modifications in “CG” context:**

~~~
minimod view --skip-supplementary hg38noAlt.fa dna.bam > minimod.tsv
modkit extract full --cpg --reference hg38noAlt.fa dna.bam modkit.tsv
awk ‘NR==1 || $14==“m”’ modkit.tsv > modkit_m.tsv
test/compare_view_mktsv_mmtsv.sh modkit_m.tsv minimod.tsv compare_out
~~~

test/compare_view_mktsv_mmtsv.sh takes modkit’s extract tsv and minimod’s view tsv and creates four files in the out_dir:

- in_both.tsv : found in both files and probability difference is <= 0.002
- large_prob_diff.tsv : found in both files and probability difference is > 0.002
- missing_in_file1.tsv : missing in file1 which is modkit_extract.tsv
- missing_in_file2.tsv : missing in file2 which is minimod_view.tsv

There are similar scripts we provide for view comparison between different file formats:

- test/compare_view_mktsv_mmtsv.sh
- test/compare_view_mktsv_mktsv.sh
- test/compare_view_mmtsv_mmtsv.sh

##### Results

compare_out/

~~~
in_both.tsv          # 721007 entries
large_prob_diff.tsv  # 0 entries
missing_in_file1.tsv # 0 entries
missing_in_file2.tsv # 0 entries
~~~

## References

1. Konstantinos Boulias and Eric Lieberman Greer. Biological roles of adenine methylation in rna. Nature Reviews Genetics, 24(3):143–160, 2023.

2. Zachary D Smith, Sara Hetzel, and Alexander Meissner. DNA methylation in mammalian development and disease. Nature Reviews Genetics, 26(1):7–30, 2025.

3. Masoumeh Fardi, Saeed Solali, and Majid Farshdousti Hagh. Epigenetic mechanisms as a new approach in cancer treatment: An updated review. Genes & Diseases, 5(4):304–311, December 2018.

4. Macsue Jacques, Kirsten Seale, Sarah Voisin, Anna Lysenko, Robin Grolaux, Bernadette Jones-Freeman, Severine Lamon, Mandhri Abeysooriya, Itamar Levinger, Carlie Bauer, et al. Meta-analysis of DNA methylation aging signatures in 17 human tissues. Nature Aging, pages 1–15, 2026.

5. Isis Beentjes, Martin A Haagmans, Desiree DSH De Bruin, Anwar Permana, Ewelina Pośpiech, Wojciech Branicki, Amade Aouatef M’charek, Kristiaan J Van Der Gaag, Titia Sijen, and Peter Henneman. DNA methylation-based forensic framework for age prediction and body fluid identification using nanopore sequencing. Forensic Science International: Genetics, page 103370, 2025.

6. Ankur Jai Sood, Coby Viner, and Michael M Hoffman. DNAmod: the DNA modification database. Journal of cheminformatics, 11(1):30, 2019.

7. Andrea Cappannini, Angana Ray, Elźbieta Purta, Sunandan Mukherjee, Pietro Boccaletto, S Naeim Moafinejad, Antony Lechner, Charles Barchet, Bruno P Klaholz, Filip Stefaniak, et al. Modomics: a database of rna modifications and related information. 2023 update. Nucleic Acids Research, 52(D1):D239–D244, 2024.

8. Lisa D Moore, Thuc Le, and Guoping Fan. DNA methylation and its basic function. Neuropsychopharmacology, 38(1):23–38, 2013.

9. Wai-Shin Yong, Fei-Man Hsu, and Pao-Yang Chen. Profiling genome-wide DNA methylation. Epigenetics & chromatin, 9(1):26, 2016.

10. Franziska Geissler, Ksenija Nesic, Olga Kondrashova, Alexander Dobrovic, Elizabeth M Swisher, Clare L Scott, and Matthew J. Wakefield. The role of aberrant DNA methylation in cancer initiation and clinical impacts. Therapeutic Advances in Medical Oncology, 16, 2024.

11. Russell P Darst, Carolina E Pardo, Lingbao Ai, Kevin D Brown, and Michael P Kladde. Bisulfite sequencing of DNA. Current protocols in molecular biology, 91(1):7–9, 2010.

12. Yang Liu, Wojciech Rosikiewicz, Ziwei Pan, Nathaniel Jillette, Ping Wang, Aziz Taghbalout, Jonathan Foox, Christopher Mason, Martin Carroll, Albert Cheng, et al. DNA methylation-calling tools for oxford nanopore sequencing: a survey and human epigenome-wide evaluation. Genome biology, 22(1):295, 2021.

13. Yilei Fu, Winston Timp, and Fritz J Sedlazeck. Computational analysis of DNA methylation from long-read sequencing. Nature Reviews Genetics, pages 1–15, 2025.

14. Anthony Rhoads and Kin Fai Au. Pacbio sequencing and its applications. Genomics, proteomics & bioinformatics, 13(5):278–289, 2015.

15. Alan Tourancheau, Edward A Mead, Xue-Song Zhang, and Gang Fang. Discovering multiple types of DNA methylation from bacteria and microbiome using nanopore sequencing. Nature methods, 18(5):491–498, 2021.

16. Onkar Kulkarni, Reuben Jacob Mathew, Rhea Jana, Lamuk Zaveri, Sreenivas Ara, Tulasi Nagabandi, Nitesh Kumar Singh, Karthik Bharadwaj Tallapaka, and Divya Tej Sowpati. Comprehensive benchmarking of tools for nanopore-based detection of DNA methylation. Nature Communications, 2026.

17. Ying Chen, Nadia M Davidson, Yuk Kei Wan, Fei Yao, Yan Su, Hasindu Gamaarachchi, Andre Sim, Harshil Patel, Hwee Meng Low, Christopher Hendra, et al. A systematic benchmark of nanopore long-read rna sequencing for transcript-level analysis in human cell lines. Nature methods, pages 1–12, 2025.

18. William Stephenson, Roham Razaghi, Steven Busan, Kevin M Weeks, Winston Timp, and Peter Smibert. Direct detection of rna modifications and structure using single-molecule nanopore sequencing. Cell genomics, 2(2), 2022.

19. Doaa Hassan, Daniel Acevedo, Swapna Vidhur Daulatabad, Quoseena Mir, and Sarath Chandra Janga. Penguin: a tool for predicting pseudouridine sites in direct rna nanopore sequencing data. Methods, 203:478–487, 2022.

20. Neda Ghohabi Esfahani, Andrew J Stein, Stuart Akeson, Talia Tzadikario, Connor Powell, Pooria Daneshvar Kakhaki, and Miten Jain. Assessment of nanopore RNA modification calling in human cell lines and synthetic systems. Genome Biology, 27(1):190, 2026.

21. Suneth Samarasinghe, Ira Deveson, and Hasindu Gamaarachchi. Realfreq: real-time base modification analysis for nanopore sequencing. Bioinformatics, 41(4):btaf151, 2025.

22. Hasindu Gamaarachchi, Hiruna Samarakoon, Sasha P Jenner, James M Ferguson, Timothy G Amos, Jillian M Hammond, Hassaan Saadat, Martin A Smith, Sri Parameswaran, and Ira W Deveson. Fast nanopore sequencing data analysis with slow5. Nature biotechnology, 40(7):1026–1029, 2022.

23. Hiruna Samarakoon, James M Ferguson, Sasha P Jenner, Timothy G Amos, Sri Parameswaran, Hasindu Gamaarachchi, and Ira W Deveson. Flexible and efficient handling of nanopore sequencing signal data with slow5tools. Genome Biology, 24(1):69, 2023.

24. Hiruna Samarakoon, James M Ferguson, Hasindu Gamaarachchi, and Ira W Deveson. Accelerated nanopore basecalling with slow5 data format. Bioinformatics, 39(6):btad352, 2023.

25. Hasindu Gamaarachchi, Sasha Jenner, Hiruna Samarakoon, James M Ferguson, and Ira W Deveson. The enduring advantages of the slow5 file format for raw nanopore sequencing data. GigaScience, 14:giaf118, 2025.

26. Heng Li. Minimap2: pairwise alignment for nucleotide sequences. Bioinformatics, 34(18):3094–3100, 2018.

27. Petr Danecek, James K Bonfield, Jennifer Liddle, John Marshall, Valeriu Ohan, Martin O Pollard, Andrew Whitwham, Thomas Keane, Shane A McCarthy, Robert M Davies, et al. Twelve years of samtools and bcftools. Gigascience, 10(2):giab008, 2021.

28. Hasindu Gamaarachchi, Chun Wai Lam, Gihan Jayatilaka, Hiruna Samarakoon, Jared T Simpson, Martin A Smith, and Sri Parameswaran. GPU accelerated adaptive banded event alignment for rapid comparative nanopore signal analysis. BMC bioinformatics, 21(1):343, 2020.

29. Jared T Simpson, Rachael E Workman, PC Zuzarte, Matei David, LJ Dursi, and Winston Timp. Detecting DNA cytosine methylation using nanopore sequencing. Nature methods, 14(4):407–410, 2017.

